# The salmon louse genome may be much larger than sequencing suggests

**DOI:** 10.1101/2022.01.14.476287

**Authors:** Grace A. Wyngaard, Rasmus Skern-Mauritzen, Ketil Malde, Rachel Prendergast, Stefano Peruzzi

## Abstract

The genome size of organisms impacts their evolution and biology and is often assumed to be characteristic of a species. Here we present the first published estimates of genome size of the ecologically and economically important ectoparasite, *Lepeophtheirus salmonis* (Copepoda, Caligidae). Four independent *L. salmonis* genome assemblies of the North Atlantic subspecies *Lepeophtheirus salmonis salmonis*, including two chromosome level assemblies, yield assemblies ranging from 665 – 790 Mbps. These genome assemblies are congruent in their findings, and appear very complete with Benchmarking Universal Single-Copy Orthologs analyses finding >92% of expected genes and transcriptome datasets routinely mapping >90% of reads. However, two cytometric techniques, flow cytometry and Feulgen image analysis densitometry, yield measurements of 1.3-1.6 Gb in the haploid genome. Interestingly, earlier cytometric measurements reported genome sizes of 939 and 567 Mbps in *L. salmonis salmonis* samples from Bay of Fundy and Norway, respectively. Available data thus suggest that the genome sizes of salmon lice are variable. Current understanding of eukaryotic genome dynamics suggests that the most likely explanation for such variability involves repetitive DNA, which for *L. salmonis* makes up ≈60% of the genome assemblies.

## Introduction

“In the future attention undoubtedly will be centered on the genome, and with greater appreciation of its significance as a highly sensitive organ of the cell, monitoring genomic activities and correcting common errors, sensing the unusual and unexpected events, and responding to them, often by restructuring the genome” – Barbara McClintock’s Nobel Lecture in 1983 [1].

A lot has been learned since 1983, and numerous genomes have been sized, sequenced and analyzed. Yet, many questions regarding genomes remain unanswered, the most fundamental potentially being: why do eukaryotic genomes vary so much in size? While complexity appears to correlate with minimum taxon genome size, the actual genome sizes bear no straightforward correlation with eukaryotic organismal complexity, even among closely related taxa, but are increasingly investigated as a trait subject to natural selection and consequently of relevance to studies of ecology and evolution [2–4]. Selective pressures in copepods have been posed for age at first reproduction in predation intense environments, resulting in smaller genome sizes [5], as well as selection for larger bodies and genome sizes in cold environments [6,7].

While genome size does not appear to govern organismal complexity, some relationships appear to be general: genome size often correlates with the proportion of noncoding, or repetitive, DNA in the genome [8,9], cell size [2,10] and growth rate [11]. Furthermore, the evolutionary importance of repetitive elements (mainly transposable elements – TEs) in lateral gene transfer [12] and generation of new phenotypes [13] is becoming increasingly apparent. This is well illustrated by TEs being responsible for more than 50% of the phenotypes emerging in *Drosophila* laboratory strains [14] and playing a role in adaptive evolution [15,16]. At the same time it must be realized that species specific effects may affect genome sizes in ways that appear to be inconsistent with the general trends: for example taxon specific allocation of phosphorus to RNA rather than nonessential non-coding DNA may result in a selection for compact genomes in phosphorus limited environments [17,18]. As the role(s) of noncoding and repetitive DNA become better understood, the importance of knowing to what extent this component of the genome has been accurately included in the assemblies and annotations becomes increasingly clear. This does not contradict the fact that partial genome assemblies may be both of high quality and immense value.

The salmon louse (*Lepeophtheirus salmonis*, Krøyer 1837) is a marine parasitic copepod of large economic and ecological importance [19,20]. It belongs to the order Siphonostomatoida and is found on salmonid fishes in the northern hemisphere. There are two *L. salmonis* subspecies separated by approximately 5 million years of evolution, *L. salmonis salmonis* (Krøyer, 1837) inhabiting the North Atlantic and *L. salmonis onchorhynchii* [21] inhabiting the Northern Pacific [21–23]. These parasites alter the physiology, disease susceptibility, growth rates, and behavior of farmed salmon [24–26] and inflict large economic losses [27]. As the salmonid aquaculture has expanded to the extent that farmed salmonids outnumber wild salmonids by 2-3 orders of magnitude in some regions in the North Atlantic, the salmon lice populations have increased in parallel and currently inflict significant economic and ecological challenges [27–28]. The combined societal and ecological impacts of *L. salmonis* have spurred intense research, including modelling to assess ecological risk [29,30], methodology for surveillance [24,31], population genetics [32–34], resistance and resilience development [35,36] and molecular biology [37–40]. As a result, the salmon louse genome has been sequenced several times using various sequencing platforms, and independent genome assemblies have been made [40,41] – including two chromosome level assemblies.

As DNA sequencing technologies have advanced and improvements in the identification and annotation of noncoding DNA have gradually followed, there is a growing awareness that genome assembly methods sometimes fail to correctly reconstruct repetitive regions and noncoding DNA [42,43]. Traditional quantitative cytogenetic methods such as flow cytometry and Feulgen microdensitometry are recognized as being reliable with respect to estimating the true amount of nuclear DNA contents and providing estimates of total genome size [44,45]. Thus, genomic assemblies and cytometric methods possess different strengths with respect to the kinds of information they provide. When these two approaches are applied in combination, they are likely to either validate the independent estimates or provide direction as to seeking explanations for the discrepancies. In light of the importance of the *L. salmonis* genome it seems prudent to use cytometric methods to validate estimates from genome sequencing based estimates.

In the present study of the salmon louse, we compare genome size estimates based on two quantitative cytometric methods (flow cytometry and Feulgen image analysis densitometry) performed on multiple samples with unpublished cytometric measurements and estimates based on whole genome sequencing. Where discrepancies are apparent, we propose explanations and a path forward for resolving them. Additionally, as this is the first published estimate of the genome size of a parasitic copepod, the genome size of this siphonostomatoid is discussed in relation to the free-living copepods in the orders Cyclopoida, Calanoida, and Harpacticoida.

## Results

### Sequencing – and assembly based genome size estimates

The salmon louse genome has been sequenced several times using various sequencing platforms, and six independent genome assemblies have been made – including two chromosome level assemblies (Table 1). The resulting assemblies appear consistent in structure, as revealed by linkage analyses [33,40,46], and sizes (Table 1), collectively suggesting the salmon louse genome size to be approximately 600-700 Mbps.

**Table 1.**
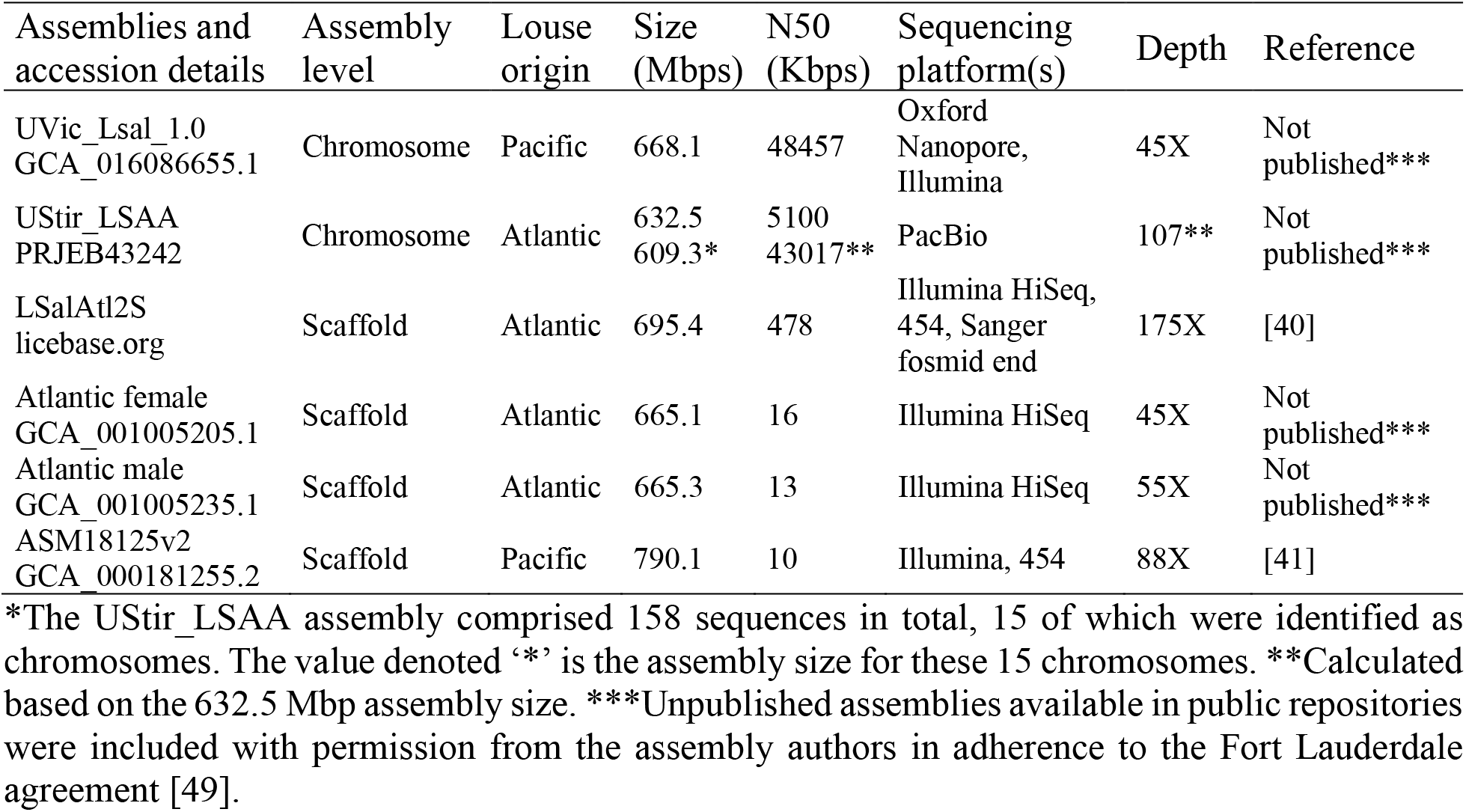
Key statistics and sequencing platforms for salmon louse assemblies available March 2021. Depths of sequencing coverage are the values associated with the assemblies and are estimated based on assembly-indicated genome sizes (i.e. ≈700 Mbps). The ASM18125v2 assembly is quite fragmented and the authors indicate that the assembly overestimates the actual size suggested to be around 600 Mbps (https://www.ncbi.nlm.nih.gov/bioproject/PRJNA40179). Only LSalAtl2s is extensively assessed and the predicted gene set appears to be quite complete as it contains 92.4% of the expected genes in a BUSCO analysis and maps ≈ 90% of transcriptome reads [47,48].

| Assemblies and accession details | Assembly level | Louse origin | Size (Mbps) | N50 (Kbps) | Sequencing platform(s) | Depth | Reference |
| --- | --- | --- | --- | --- | --- | --- | --- |
| UVic_Lsal_1.0<br>GCA_016086655.1 | Chromosome | Pacific | 668.1 | 48457 | Oxford Nanopore, Illumina | 45X | Not published*** |
| UStir_LSAA<br>PRJEB43242 | Chromosome | Atlantic | 632.5<br>609.3* | 5100<br>43017** | PacBio | 107** | Not published*** |
| LSalAtl2S<br>licebase.org | Scaffold | Atlantic | 695.4 | 478 | Illumina HiSeq, 454, Sanger fosmid end | 175X | [40] |
| Atlantic female<br>GCA_001005205.1 | Scaffold | Atlantic | 665.1 | 16 | Illumina HiSeq | 45X | Not published*** |
| Atlantic male<br>GCA_001005235.1 | Scaffold | Atlantic | 665.3 | 13 | Illumina HiSeq | 55X | Not published*** |
| ASM18125v2<br>GCA_000181255.2 | Scaffold | Pacific | 790.1 | 10 | Illumina, 454 | 88X | [41] |

\*The UStir\_LSAA assembly comprised 158 sequences in total, 15 of which were identified as chromosomes. The value denoted ‘\*’ is the assembly size for these 15 chromosomes. \*\*Calculated based on the 632.5 Mbp assembly size. \*\*\*Unpublished assemblies available in public repositories were included with permission from the assembly authors in adherence to the Fort Lauderdale agreement [49].

To evaluate the congruence of the assemblies, while not requiring them to conserve synteny, we created libraries of 240 bp synthetic reads from each of the assemblies. These synthetic reads were then mapped to each of the published assemblies using BLAST. The results show that the assemblies are close to interchangeable in terms of their sequence composition (Table 2) and that the differences in the sequence captured by the different sequencing technologies are minor.

**Table 2.**
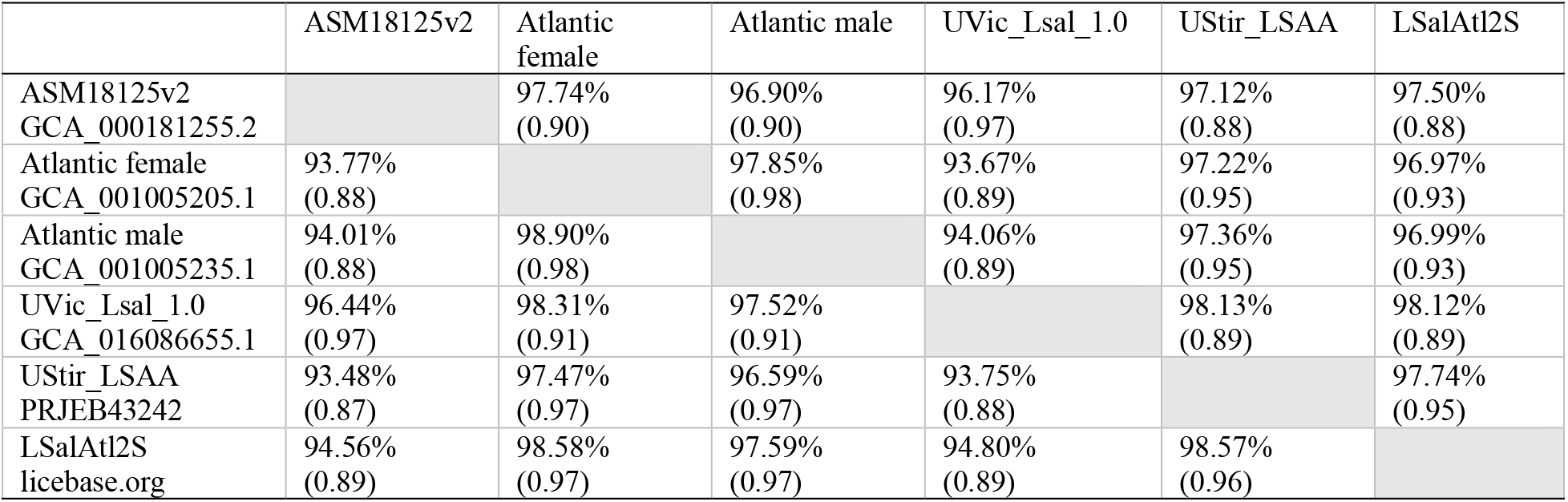
Consistency of assemblies. The assemblies were converted to 240 bp synthetic reads that were blasted against all other assemblies. The assemblies from which synthetic reads originated are indicated in column headings and the reference assemblies are indicated in the row headings. The results show average query cover % and (the proportion of reads that maps with >95% identity).

It is well known that genome assembly sizes may deviate significantly from the actual genome size [9] and additional sequence-based genome size estimates were therefore produced. First, k-mer analysis was performed using Jellyfish [50]. The published GLW4 dataset originating from inbred salmon lice [40] and word sizes (k-mer lengths) of 21 – 31 yielded estimated genome sizes of 976 – 1017 Mbps (modal k-mer coverages: 20 – 17). Repeating the analysis using previously published data from wild salmon lice [33] and using the word lengths of 29 and 31 yielded estimated genome sizes of 1086 and 1015 Mbps (modal k-mer coverages: 55 and 53).

Second, sequencing reads were mapped to the LSalAtl2s genome using BWA [51]. For each library, modal coverage (M) was extracted, and assumed to be representative of diploid coverage. All coverages for the genome were then summed and divided by the modal coverage to estimate the genome size. Under the assumption that repeat sequence occurring N times in the genome would have a coverage distribution centered on N*M, each location in the repeat would be counted N times. Seven inbred salmon lice libraries [40] were used, with modal coverages of 3–23x, and resulting in genome size estimates of 791-1184 Mbps, with low coverage libraries giving the highest size estimates. The analysis was repeated with six libraries from wild lice [33], giving coverages of 5–17x, and size estimates of 845–1073, again with low coverage libraries leading to higher size estimates.

### Genome size estimates based on FCM

The nuclear DNA contents of somatic (2*C* values) and gametic cells measured by flow cytometry (FCM, Runs 1–5) in gametic, naupliar or adult stages of Atlantic *L. salmonis salmonis* are reported in Table 3. Figure 1 shows representative fluorescence (FL histograms of the above cells analyzed together with reference standards). Overall, fluorescence histograms of propidium iodide (PI) - stained somatic cells or gametes indicated good resolution levels with coefficients of variation (CVs) in the range of 1-5% (Table 3).

**Table 3.**
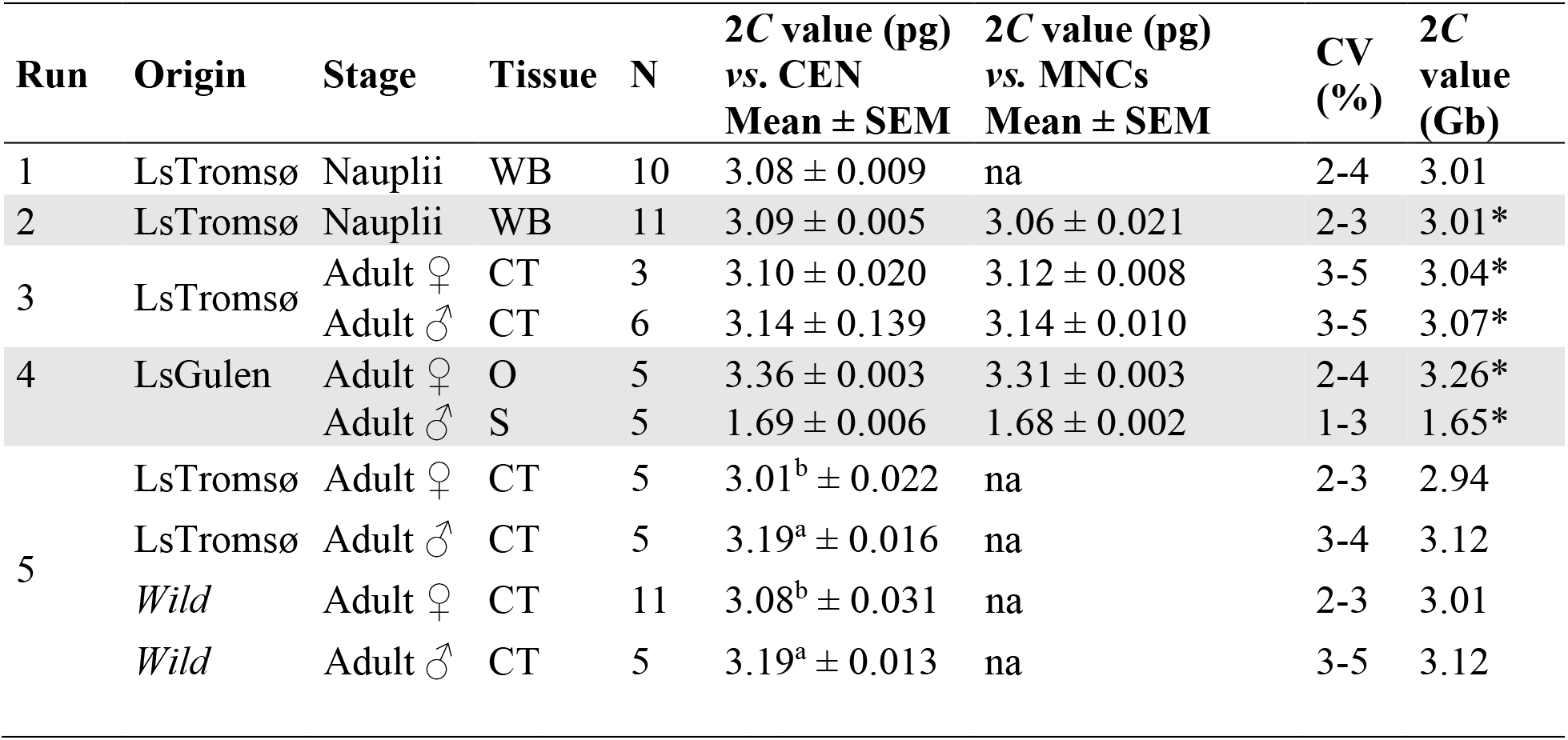
Nuclear DNA contents of somatic and gametic cells of *L. salmonis salmonis* as measured using flow cytometry. Runs 1, 2, 3, and 5 measured somatic cells of nauplii or adults of the LsTromsø strain. Run 4 measured gametic cells of the *Ls* Gulen laboratory strain. Run 5 compared the laboratory reared LsTromsø strain to wild caught adults from naturally infected fish reared in Tromsø. Chicken and/or human white blood cells were used as internal reference standards. CEN = Chicken erythrocyte nuclei (2*C* value = 2.5 pg DNA per nucleus), MNCs = human mono-nucleated cells (2C value = 7.0 pg DNA per nucleus), WB = whole body, CT = cuticular and subcuticular tissues from cephalic region, O = oocytes, S = sperm, N = number of individuals or number of samples in the cases of nauplii analyzed, CV= Coefficient of Variation as a percentage of mean for target nuclei (*L. salmonis* data). Na = not available. For naupliar stages (Run 1 and 2), each sample consisted of approximately 50 nauplii. *Average value based on the two internal standards. For Run 5, 2C values with superscripts “a” and “b” differ at P < 0.05 (two-way ANOVA).

**Figure 1.**
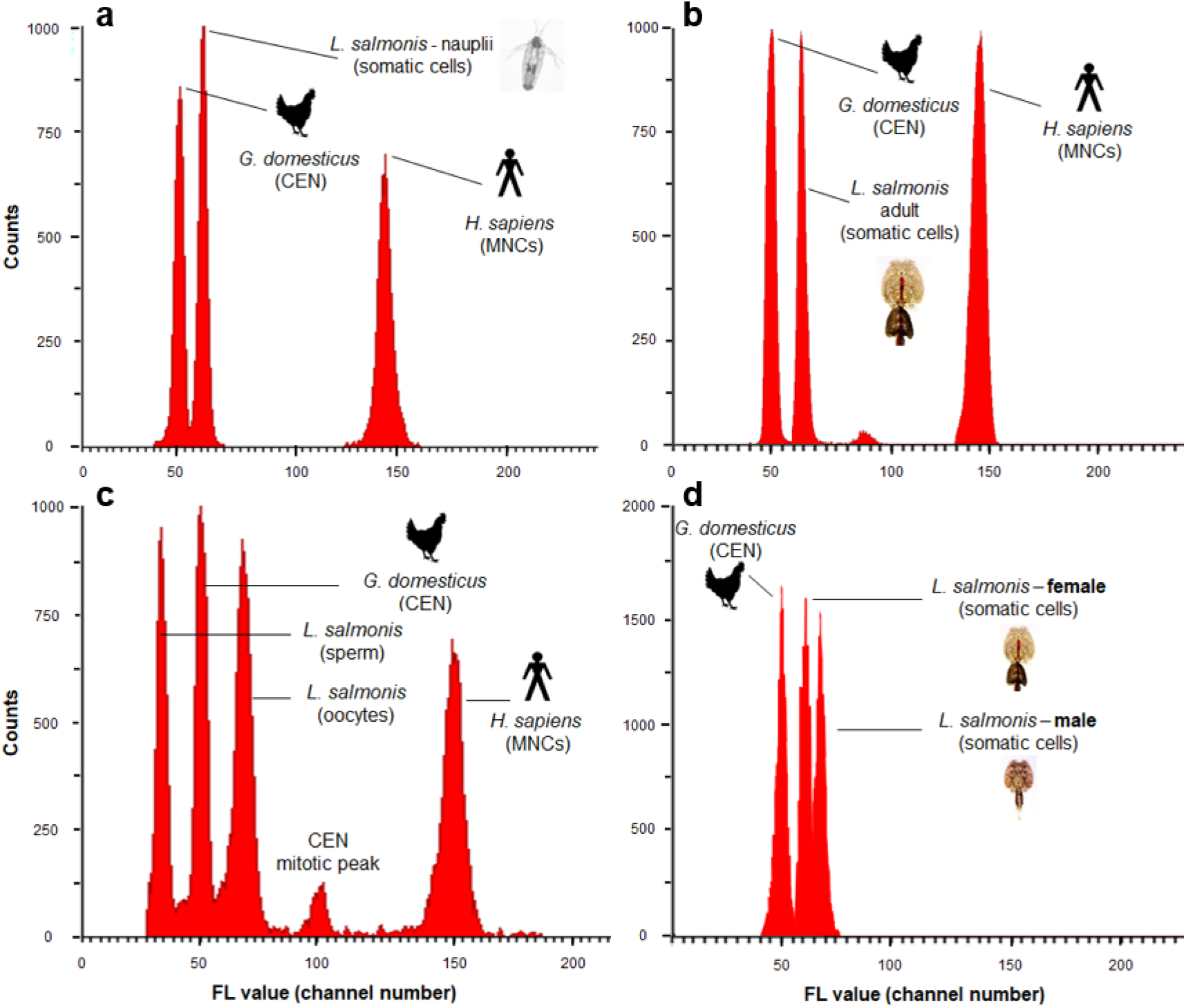
Representative fluorescence histograms of PI-stained somatic and gametic cells of *L. salmonis salmonis*. (a) Run 1, somatic cells obtained from whole body squash of nauplii; (b) Run 3, somatic cells obtained from cephalic regions of adult females; (c) Run 4; oocyte and sperm samples; (d) Run 5, somatic cells obtained from cephalic regions of adult males and females. Samples analyzed using chicken and human cells as internal reference standards. Fluorescence (FL) peaks values on the X-axis are reported in arbitrary units expressed as Propidium Iodide (PI) fluorescence channel numbers (FL value). Values on the Y-axis (counts) refer to the number of nuclei counted per channel. CEN = chicken erythrocytes nuclei, MNCs = human mono-nucleated cells.

The estimated 2*C* DNA contents of somatic cells in naupliar stages of Ls Tromsø were similar across replicates (FCM Runs 1 and 2), averaging 3.08 pg DNA per nucleus when using chicken (CEN) as an internal standard (Table 2) and did not vary significantly when both chicken and human (MNCs) standards were used simultaneously (FCM Run 2). Nauplii cannot visually be assigned a sex and thus it cannot be known for certain what sex ratio occurred in these samples although sex is genetically determined with a 50:50 ratio [40].

The average nuclear 1*C* DNA contents of Ls Gulen sperm cells (FCM Run 4) were 1.68 and 1.69 pg DNA per nucleus, and did not differ when estimated using chicken or human standards. The estimated 2*C* value of oocytes (FCM Run 4) averaged 3.33 pg DNA per nucleus with no significant difference between values based on the two standards. A derived 2*C* value of sperm cells (twice the 1.69 pg DNA per sperm cell) does not differ significantly from that of the unfertilized oocytes, 3.33 pg DNA per nucleus (Table 2).

The nuclear DNA contents of males and females in both wild caught and laboratory reared Tromsø strains were compared in Run 5. The 2*C* DNA content of somatic cells averaged 3.01 and 3.19 pg DNA/nucleus in female and male specimens of a laboratory strain, respectively. The same trend was observed when analyzing somatic cells of wild specimens; the 2*C* values averaging 3.08 and 3.19 pg DNA per nucleus in females and males, respectively. Overall, within this FCM run male genome size estimates were consistently larger (ANOVA, P=0.0001) that female genome size estimates whereas, the 2*C* DNA contents of somatic cells within a gender did not differ significantly between salmon lice of different origin and with no interaction between the two factors. Despite the lack of a statistically significant difference between male and female adults of laboratory reared Ls Tromsø salmon lice recorded in FCM Run 3, the DNA content of these somatic cells did not significantly differ from the laboratory and wild caught adults measured in Run 5, when comparisons were made within gender.

### Feulgen image analysis densitometry (FIAD) – Nuclear morphologies of somatic tissues

The squashed somatic tissues possessed a variety of nuclear morphologies, from dense to diffuse (Fig. 2), which yielded a corresponding variety of values of integrated optical density (IOD). Such heterogeneity in nuclear morphology was not observed in either chicken or trout erythrocyte standards (Fig. 2a,b). We made decisions to select for measurement nuclei with intermediate morphologies that possessed a granular and slightly diffuse appearance (Fig. 2c,d). Nuclei that were very densely staining in appearance, or compact (Fig. 2e), yielded IOD values at the lower end of the range, possibly due to DNA compaction. Nuclei with a very diffuse appearance and sometimes with nuclear membranes possessing uneven edges that might indicate partial degradation (Fig. 2f), tended to possess IOD values at the higher end of the range. The corresponding 2*C* values were 2.1 pg DNA per nucleus for densely stained nuclei to 3.4 pg DNA per nucleus for diffuse nuclei.

**Figure 2.**
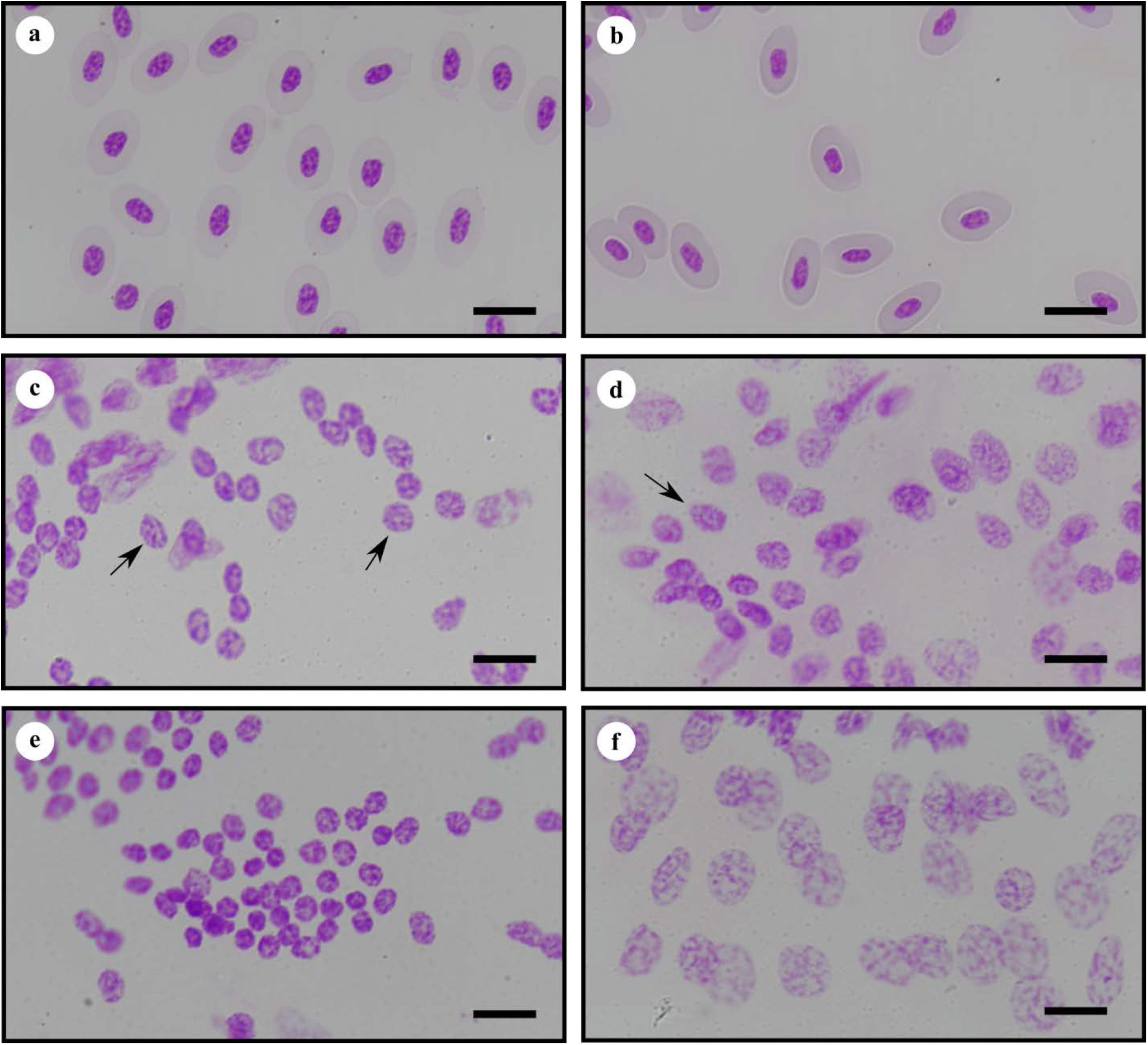
Photomicrographs of squash preparations of somatic nuclei of cephalothorax of *L. salmonis salmonis* and erythrocytes of hen and trout used as standards, stained with Feulgen reaction for DNA. (a) Nuclei of erythrocytes of *G. domesticus;* (b) Nuclei of erythrocytes of *O. mykiss;* (c) Representative nuclei, indicated by arrows, of male *L. salmonis salmonis* slide 64 whose measurements are given in Table 4; (d) Representative nucleus, indicated by arrow, of female *L. salmonis salmonis* slide 63 whose measurements are given in in Table 4; (e) Nuclei of male *L. salmonis salmonis* slide 64 with high level of DNA compaction; (f) Nuclei of male *L. salmonis salmonis* slide 64 with highly diffuse morphology. Scale bars represent 10 μm.

### Genome sizes based on FIAD

Based on Feulgen image analysis densitometry of the Ls Gulen laboratory strain and the chicken standard, the average somatic nuclear DNA contents of individual adult males (2.86 – 2.93 pg DNA per nucleus) were consistently larger than the average nuclear DNA contents of individual females (2.64 – 2.78 pg DNA per nucleus) (Table 4). Despite the consistency of the trend, the average of three individual females, 2.70 pg DNA per nucleus, did not differ significantly from the average of three individual males, 2.90 pg of DNA per nucleus (Table 4). The two sample *t* tests were based on 3 individuals per sex, rather than 120 nuclei per sex (Table 4; the N values for LsGulen), to avoid pseudoreplication.

**Table 4.** Nuclear DNA contents of somatic cells of individual adult *L. salmonis* as measured using Feulgen image analysis densitometry. Cephalothorax (CT) tissues of laboratory reared adults were obtained from the *Ls* Gulen and *Ls*1a populations. Tissue from the spine of an appendage (SP) was obtained from a wild caught Maine population. Hen (CEN) and male human mononucleated leucocytes (MNCs) were used as internal reference standards to estimate values in picograms (pg). Values based on hen were converted to gigabases (Gb). N refers to number of nuclei measured in each adult. SEM refers to standard error of the mean; CV refers to coefficient of variation of IOD values.

| Origin | Stage | Tissue | N | 2C value (pg)<br>vs. CEN<br>Mean ± SEM | 2C value (pg)<br>vs. MNCs<br>Mean ± SEM | CV<br>(%) | 2C<br>value<br>(Gb) |
| --- | --- | --- | --- | --- | --- | --- | --- |
| *Laboratory* |  |  |  |  |  |  |  |
| *Ls* Gulen 61 | Adult ♀ | CT | 13 | 2.78 ± 0.012 | na | 1.6 | 2.74 |
| *Ls* Gulen 62 | Adult ♀ | CT | 15 | 2.64 ± 0.003 | na | 3.3 | 2.60 |
| *Ls* Gulen 63 | Adult ♀ | CT | 92 | 2.67 ± 0.003 | na | 10.4 | 2.63 |
| Average |  |  |  | 2.70 ± 0.043 | na |  | 2.66 |
| *Ls1a* 133 | Adult ♀ | CT | 34 | 2.64 ± 0.046 | 2.60 ± 0.046 | 10.2 | 2.60 |
| *Ls1a* 134 | Adult ♀ | CT | 20 | 2.75 ± 0.037 | 2.71 ± 0.037 | 6.1 | 2.67 |
| *Ls1a* 137 | Adult ♀ | CT | 16 | 2.71 ± 0.025 | 2.67 ± 0.025 | 3.9 | 2.65 |
| Average |  |  |  | 2.70 ± 0.032 | 2.66 ± 0.032 |  | 2.66 |
| *Ls* Gulen 64 | Adult ♂ | CT | 65 | 2.93 ± 0.035 | na | 9.9 | 2.89 |
| *Ls* Gulen 65 | Adult ♂ | CT | 36 | 2.86 ± 0.037 | na | 7.9 | 2.82 |
| *Ls* Gulen 66 | Adult ♂ | CT | 19 | 2.90 ± 0.054 | na | 8.3 | 2.85 |
| Average |  |  |  | 2.90 ± 0.020 | na |  | 2.85 |
| *Wild* |  |  |  |  |  |  |  |
| Copscook Bay, Maine | Adult ♀ | SP | 3 | 3.07 ± 0.176 | na | 9.8 | 3.00 |

A single wild caught adult female collected from a Maine population possessed nuclei in an appendage that were especially well suited for measurement as they were well isolated and lacked any visible background stain. The mean value of these nuclei, 3.07 pg DNA per nucleus, does not differ from the wild female that was caught near Tromsø nor from the laboratory reared Ls Tromsø strain (Tables 3, 4). It should be noted that the Maine specimen was squashed and stained as above, but prior to employing the freeze cracking technique, and thus measurements were restricted to those few nuclei in the spine of a swimming leg that possessed a granular and slightly diffuse appearance, and were also isolated and surrounded by clear background.

### Comparison of genome size estimates based on FIAD and FCM

Estimated 2*C* genome sizes of male and female laboratory reared *Ls* Gulen adults obtained using FIAD and based on the chicken standard (2.90 and 2.70 pg DNA per nucleus, respectively, Table 4) are within 10% of the estimates obtained using FCM on the Ls Tromsø laboratory strain (Table 3), assuming values of 3.2 and 3.0 pg DNA per nucleus for males and females, respectively, for the Ls Tromsø strain). Each slide containing a population of nuclei from a single adult in the FIAD analyses contained some values that overlapped with estimates obtained using FCM; however, these higher FIAD values did not equal the central tendency of values obtained using FCM. The average nuclear DNA content obtained for the Ls Gulen adult females using FIAD, 2.70 pg DNA per nucleus, is within the 15% range of the value based on oocytes 3.26 pg DNA per nucleus (FCM run 4).

## Discussion

*L, salmonis* assemblies are consistently approximately 700 Mbps suggesting that the *L. salmonis* 1*C* genome may be of approximately the same size, or possibly larger if the assemblies include substantial collapsed repeated regions. This approximate size was readily accepted by the salmon louse research community, as the first cytometric based estimate of a North Atlantic population was 0.58 pg (≈ 567 Mbps) [52]. Thirteen years later a Bay of Fundy population measured in the same lab was estimated to be 0.96 pg (≈ 939 Mbps) [53]. Yet in the present study the *L. salmonis* subsp. *salmonis* genome size estimates range from 1.3 to 1.6 Gbps when determined by two independent cytometric methods being applied to three laboratory strains and wild salmon lice from two locations. Sequence based extrapolations, in contrast, yield estimates ranging between 0.8 and 1.2 Gbps. In attempting to identify the factors responsible for the different estimates of *L. salmonis salmonis* genome size, we emphasize the importance of harnessing complementary approaches to estimate genome size and where discrepancies exist, not to disregard potentially ‘missing’ portions of DNA which may play an important role in adaptation. Additionally, while it is a common bias to interpret ‘old measurements’ as wrong when they disagree with new measurements, we remain open to a plethora of explanations to reconcile older and present study findings based on cytometric methods.

Genome assembly sizes are known to be unreliable predictors for genome size. Highly conserved repeats in combination with read errors can be difficult to resolve, and assemblies can have repeated regions collapsed into one sequence or have multiple copies of the same genomic region. More precise estimates can be achieved by examining the sequence reads. We have used two approaches, one using k-mer statistics and another based on mapping statistics against a reference assembly. These methods point to a genome size of 800-1200 based on mapping, and 1000-1100 Mbp based on k-mers. One partial explanation for the variability can be unmapped reads (approximately 5% of the reads). If they represent sequence not present in the genome assembly, these genome components will be omitted from the mapping estimate, but included in the k-mer estimate. In addition, most of the sequence data is from female salmon lice, and both approaches will count the average of the haploid sex chromosomes, not their sum.

Flow cytometry is a well-established method for nuclear DNA content analysis and characterization in experimental biology, and is increasingly being used due to its rapidity, precision, and reproducibility. Feulgen image analysis densitometry similarly has experienced a resurgence in its use, partially due to its affordability and applicability when the number of available nuclei to measure is small. Interpretations of genome size estimates based on FCM and FIAD require careful consideration of the advantages and disadvantages of each method for the species and specific tissues under consideration. Explanations of differences among cytometric measurements in general, as well as those in the present study with unpublished estimates [52,53], performed on *L. salmonis* could include tissue compaction levels that can cause estimates to vary two-fold, misidentification of haploid and diploid cells, and misidentification of species, the latter of which seems quite unlikely as species identifications were provided by external expert collaborators. Not to be discounted is the possibility of real variation in genome size among populations. Jeffrey’s [53] review highlights the issues of concern, particularly the chemistries and tissues measured most applicable to the present study, and are discussed in Supplemental Methods.

Measurements of nuclear DNA contents of naupliar and/or adult stages of two laboratory strains and wild caught *L. salmonis salmonis* based on FCM and two laboratory strains and one wild caught population of adults stages based on FIAD estimate *2C* somatic nuclear contents to be 3 ± 0.3 Gb. Halving the FCM values to obtain the 1*C* amount and directly measuring sperm DNA content yielded values ranging from 1.47 – 1.65 Gb (Table 3). Gametic nuclei of *L. salmonis salmonis* contained one half the DNA of the somatic nuclei, and indicates a lack of chromatin diminution, a phenomenon in which 1 *C* values cannot be estimated by halving somatic genome sizes [54].

Male genome size was consistently slightly larger than female genome size, expectedly so due to erosion of the W-chromosome in the heterozygotic female [55]. There was also no evidence of mitoses in the adult somatic tissues, and therefore most adult somatic cells are suitable for genome size measurement [56]. Furthermore, we found no evidence of significant differences based on cytometric based comparisons of geographical (Norway, Maine), laboratory (Ls1a, Ls Gulen and Ls Tromsø) or wild caught (Norway, Maine) populations.

Genome size estimates of crustaceans based on FIAD are commonly lower than those based on FCM and estimates within 15% of one another are generally considered reliable [53]. Accordingly, the FIAD derived estimates of 2.7 and 2.9 pg DNA per nucleus for adult females and males respectively, are within 10% to 15% lower than the FCM based estimates, depending upon the particular comparison. The most likely explanation for the disparity between FCM and FIAD based estimates is that the laser detection of cells in a suspension used in FCM is less sensitive to background noise, DNA compaction and other conformational changes in the chromatin sometimes encountered when measurements are based on the quantitative intercalation of the Schiff reagent among nucleotides as they sometimes are in squashed tissues in FIAD. These differences between FCM and FIAD based estimates of adults correspond to approximately 0.3 pg in a 2C nucleus, or ≈ 150 million base pairs in the 1*C* genome. We conclude that measurements obtained from FIAD and FCM were internally consistent and that the discrepancy between the results are well within the boundaries expected from earlier studies [57]. Since cytometric measurements are based on direct observations and the derived estimates are both consistent within the methods and discrepancies in accordance with methodological expectations we regard the cytometric results, collectively indicating a *L. salmonis salmonis* genome size of approximately 1.5 Gbps, as the most reliable measurements available.

The above suggests that sequence based methods underestimate the genome size by approximately 1/3^rd^. The mapping approach is sensitive to errors in assembly completeness, uniformity of library and sequencing coverage, mapping accuracy and modal mapping estimation. Alas, our analysis did not reveal which of these factors were the more likely to cause the apparent mapping based underestimation of genome size. As an alternative to the mapping based estimates we applied the widely used k-mer approach which similarly appeared to miss approximately 1/3^rd^ of the genome size. The most plausible explanation may be that repetitive elements cause the k-mer approach to underestimate genome size as previously observed by Pflug and co-workers [58]. Similarly, genome assembly based on k-mer analysis of the lobster *Homarus americanus* is believed to be missing approximately 28% of the genome [59]. The salmon louse genome is among the crustaceans with highest occurrence of repetitive elements; ≈60% of the assembly annotated as repeats [40], suggesting that such underestimation of size may not be implausible. In the present study sequence based genome size estimates consistently provided substantially lower genome size estimates than cytometric measurements. Hence estimates should be regarded with caution until they have been confirmed by direct cytological measurements.

While our cytometric based estimates converge on a genome size of approximately 1.5 Gbps, earlier cytometric measurements disagree: a 1*C* = 0.58 pg DNA per cell (567 Mbp) estimate by Gregory [52] was based on material of Norwegian *L. salmonis salmonis* from a discontinued lab strain supplied by Professor Frank Nilsen of the University of Bergen in the early 2000’s (pers. comm. Frank Nilsen) and a later 1*C* = 0.96 pg DNA per cell (939 Mbp) estimate by Jeffery [53] which was based on material collected in the Bay of Fundy in the 2010’s and supplied by Professor Elizabeth Boulding from the University of Guelph. The most parsimonious explanation of the discrepancy is to consider the earlier measurements erroneous. However, the measurements were made by respected authorities in the field, with whom our measurements on other species have agreed, and the salmon lice were supplied by competent scientists. We therefore believe their measurements are unlikely to be incorrect and consider the results to indicate that large variations in the salmon louse genome size occasionally arise. Such variability would not be unprecedented in copepods. Based on FCM geographically based intraspecific variations in genome size of magnitudes 1 – 9 pg have been reported in the marine calanoid copepods *Calanus glacialis*, *Calanus hyperboreus*, and *Paraeuchaeta norvegica* populations inhabiting the High Arctic and Southern fjords of Norway [6]. Based on FIAD a difference of 1 pg between German (Schöhsee “house” lake of current Max Planck Institute for Evolutionary Biology in Ploen) and Lake Baikal populations of the freshwater cyclopoid *Mesocyclops leuckarti* was reported [60,61]. Furthermore, the genome size of the North Sea population of the calanoid *Pseudocalanus elongatus* decreased after being reared in the laboratory for 96 generations [62]. It would not be surprising to encounter additional examples with intraspecific genome size differences in other copepods.

The somatic 2*C* nuclear DNA contents of *L. salmonis*, as estimated using FCM and FIAD in laboratory and wild populations (2.7 – 3.2 pg DNA per nucleus and corresponding to genome size of 1.33 – 1.56 Gbps) are at the lower end of the range of all published values of free-living copepods, which vary more than 300 fold from 0.20 – 64.46 pg DNA per nucleus, corresponding to genome sizes ranging from 195 Mbps to 63 Gbps (Fig. 3). Relative to the range of values in free-living cyclopoids, *L. salmonis salmonis* has an intermediate genome size, although its genome size is larger than the majority of cyclopoid species estimates. Relative to the range of cytometric based values in calanoids, *L. salmonis salmonis* is comparable to the smaller genomes, with the majority of calanoid species possessing larger or far larger genomes.

**Figure 3.**
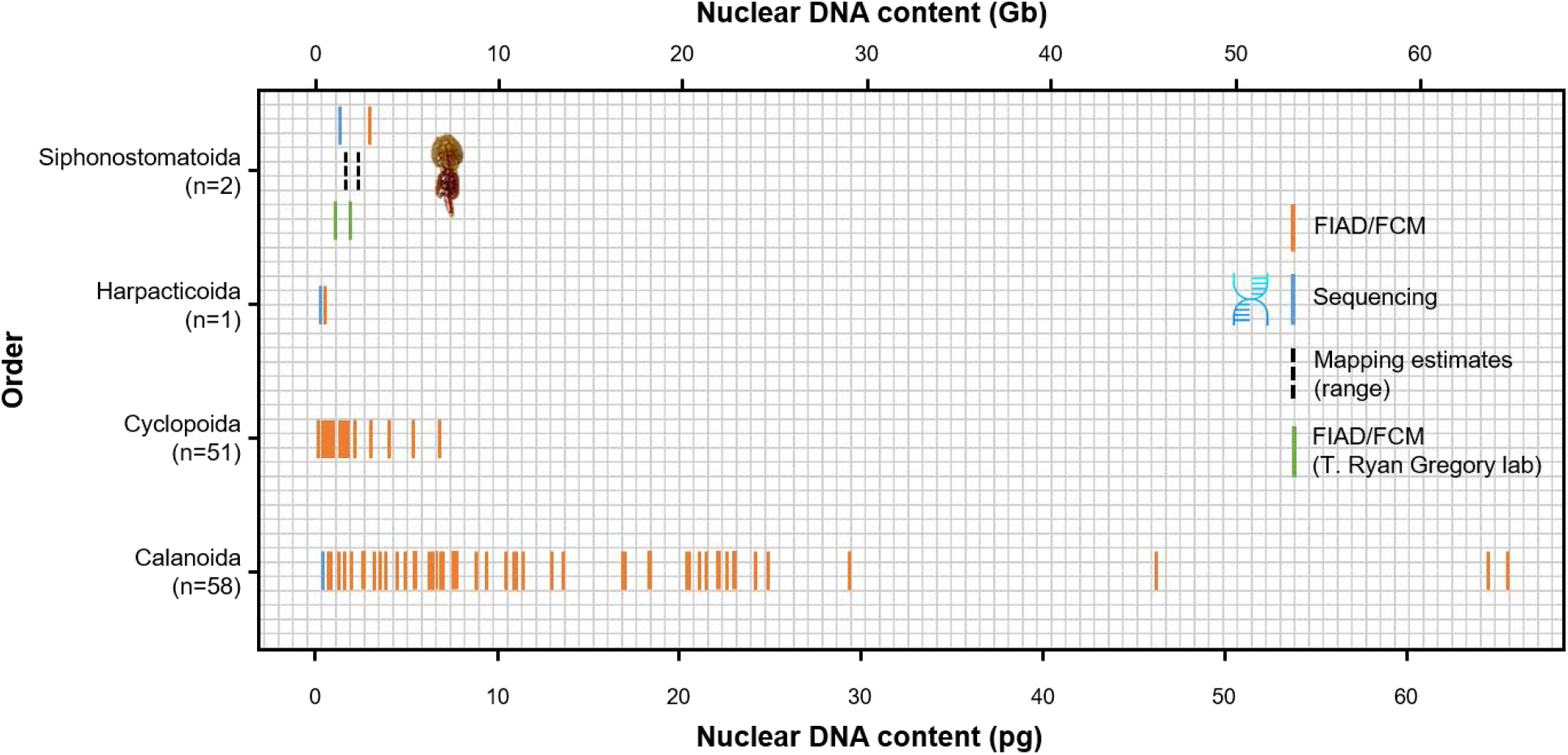
Somatic nuclear DNA contents of copepods. Each species is represented by a vertical bar. Estimates based on cytometric methods, genome assemblies, and mapping are presented in orange, blue, and dotted black, respectively, for the siphonostomatoid *L. salmonis salmonis*. The black dotted lines depict a range of values obtained for *L. salmonis salmonis* using mapping. FIAD and FCM measurements depicted by green vertical lines for *L. salmonis salmonis* are from Gregory [52] and Jeffery [53], respectively. FIAD (orange) and genome assembly (blue) estimates are shown for the harpacticoid *T. californicus* [63,64]. FIAD (orange) and genome assembly (blue) estimates for the calanoid *E. affinis* are the two left most vertical lines in the distribution of Calanoida [65,66]. The adult somatic genome size estimates of Cyclopoida and Calanoida (orange) were mostly acquired using FIAD and are from Gregory [52]. All estimates are presented as 2*C* values found in the soma because some cyclopoid species and at least one calanoid species possess embryonic chromatin diminution, in which germline genome sizes (conventionally noted as 1 *C*) substantially exceed the sizes of somatic values, thus violating the appropriateness of halving 2*C* estimates in the soma to obtain 1*C* estimates of the germline.

We speculate that the high abundance of repetitive regions in the *L. salmonis salmonis* genome facilitates the observed variability in genome size by serving as a “size accordion” [67], and further suggest that a lower limit to the genome size likely exists, in a manner similar to what is indicated for the rotifer *Brachionus asplanchnoidis* in which genome size varies by a factor of 1.9 [68]. A consequence of a possible variability in genome size is that the mapping and k-mer based size estimates should not be considered conclusively discredited as they are based on sequence information from different wild samples or strains. For the Ls1a strain that was measured both by FIAD, FCM and included in the mapping and k-mer based analysis, the discrepancy may be assumed to be genuine, although the samples for cytometric analyses were sampled more than 10 generations after the samples for sequencing were obtained. Hence the *L*. *salmonis salmonis* LSalAtl2s genome assembly [40] appears to fail to resolve ≈50% of the genome that is captured in the cytometric measurements. There are several possible sources that may contribute to this discrepancy: failure to isolate some regions of DNA, failure to capture all DNA in sequencing libraries or sequencing reactions, errors introduced during the bioinformatic analyses and specific challenges caused by TE’s and DNA repeats.

The fact that the six independent genome assemblies are congruent in content despite originating from DNA purified from multiple origins of *L. salmonis* in different laboratories and sequenced using various sequencing platforms (Illumina, 454 pyrosequencing, Oxford nanopore, PacBio and Sanger) may suggest that no parts of the genome are systematically missed by the purification and sequencing protocols applied. However, systematic omission of genetic regions across isolation protocols or biased representations of repetitive regions cannot be excluded and would yield biased or incomplete genome representations. A similar distorting effect may be introduced during downstream bioinformatic analysis, for instance, by collapsing repetitive regions [43]. Distinguishing between a bias in sequencing representation and bioinformatic artifacts as the cause of underestimating repetitive regions is challenging. PacBio and Nanopore sequencing platforms produce long reads that are commonly suggested as a tool to address repetitive regions. However, the two most recent *L. salmonis salmonis* assemblies were produced using PacBio (UStir_LSAA, Table 1) and Oxford nanopore combined with Illumina (UVic_Lsal_1.0, Table 1) sequencing. These did not deviate significantly in content or size from earlier assemblies and hence did not resolve the question of the missing DNA. Since there are no indications that specific regions are missed in any of the assemblies (Table 2) it is possible that the majority of the ≈ 800 Mbp of “missing” DNA (cytometric based genome size minus genome assembly size) in the samples measured by FIAD and FCM in the present study is comprised of TEs and DNA repeats that are not accurately captured in the assemblies. The fact that the long read based assemblies do not, at least partly, resolve the challenge may indicate that mini- and microsatellites may be dominating the missing fraction since these are considered more prone to incorrect rendering by long read sequencing than TE’s [43,69]. Hence, the ≈1500 Mbp genomes measured in the present study suggestively consist of ≈ 80% (≈ 1200 Mbp) repetitive or otherwise uncaptured regions and ≈20% (≈300 Mbp) non-repetitive regions.

Challenges in capturing repeated regions are likely exacerbated in mid to large size genomes. At the lower range of genome sizes in copepods are the tidepool harpacticoid copepod *Tigriopus californicus* and the estuarine calanoid copepod *Eurytemora affinis* whose 2*C* genome sizes estimated by FIAD are 0.5 and 0.6–0.7 pg DNA per nucleus, respectively [63,65]. Both genome assemblies were significantly smaller by ≈20% (~ 400 Mb for *T. californicus*, ~ 495 Mb for *E. affinis* for 2*C* values) which attributed to the inability to sequence all of the repetitive DNA [64,66] (Fig. 3). While *L. salmonis salmonis* has a genome size at the lower range of the distribution of copepods (Fig. 3), it is still two to three fold larger than *T. californicus* and *E. affinis*. A significant proportion of repetitive regions of the salmon louse genome consists of transposable elements, or of unclassified repeated motifs that may in time be annotated as TEs [40]. Precisely identifying the portion of the genome that is comprised of TEs and their composition is of interest as TEs are increasingly viewed as drivers of genome plasticity that facilitate the rise of new phenotypes, such as acquiring insecticide resistance in fruitflies [70,71]. It may be speculated that the high occurrence of repeated regions, including TEs, in the salmon louse genome may have contributed to its documented ability to develop resistance towards new medicinal treatments despite a low diversity of genes typically associated with detoxification and stress response [36,40,72–74]. If this is the case, the use of medicines in salmon farms that harbor the majority of sea lice in parts of the North Atlantic may have positively selected for high numbers of TEs and hence for a larger genome size. This hypothesis is directly testable as *L. salmonis onchorhynchi* in the Pacific is less influenced by salmon farming, owing to the large stocks of wild salmonids, and hence should display a more frequent occurrence of smaller genomes.

## Materials and Methods

### Assembly analyses and sequencing-based genome size estimates

To evaluate the congruence of the six assemblies available in public databases (Table 1), while not requiring them to conserve synteny, we made a script that converted the individual assemblies into non-overlapping 240 bp synthetic reads thus generating 6 synthetic read libraries. These synthetic reads were then mapped to each of the published assemblies using BLAST with the following command line: *blastn -num threads 16 -evalue 1e-10 -outfmt 6 -num alignments 10 -penalty -1 - reward 1 -gapopen 3 -gapextend 2*. Since the same fragment may be mapped multiple times we calculated the percentage of synthetic reads that mapped and the fraction of the mapped reads that mapped with >95% identity (Table 2).

The genome size was estimated from sequencing and assembly data using three different approaches: Modal mapping extrapolation, k-mer analysis and single copy gene mapping extrapolation. The assembly based estimates were derived using the LSalAtl2s assembly [40]. The LSalAtl2s assembly was compared to other available assemblies (Table 1) to reveal potential regions that are missing.

Modal mapping extrapolation is based on the assumptions that populations of non-repetitive DNA sequence reads follow a Poisson distribution and this makes up the majority of DNA. By finding the modal coverage and dividing the total number of sequenced bases mapped by this number, we can estimate the genome size. This is done using the Lande-Waterman formula; G = NL/C, where G is the genome size, N is the number of reads, L is the average read length, and C is the modal coverage. The modal coverage was determined by plotting the number of sites against nucleotide coverage and identifying the peak value. To facilitate this, sequence reads from the GLW4, GLW13 and GLW16 libraries [40] previously published were mapped against the LSalAtl2S genome assembly using Samtools [51].

K-mer analysis is based on the assumption that the possible words of a certain size in a genome (k-mers) increase with the size of the genome. A genome consisting primarily of non-repetitive DNA regions will generate an approximately random population of k-mers, and the diversity of k-mers in a population of reads can be used to estimate the genome size. In the present study k-mer analyses were performed using Jellyfish [50] and sequence reads from the GLW4 library derived from a laboratory strain of *L. salmonis salmonis* [40] and libraries derived from *L. salmonis salmonis* collected in the field.

### Nuclear DNA content analysis by flow-cytometry (FCM)

#### Field and laboratory populations

Specimens of *L. salmonis salmonis* were obtained from several sources: (1) Wild adult males and females were collected from naturally infected farmed Atlantic salmon held at the sea cage facilities of the Aquaculture Research Station in Tromsø (FCM Run 5); (2) An outbred laboratory strain, *Ls* Gulen, was derived from adults collected in *Ls* Gulen (Norway) and reared at the Salmon Louse Research Centre in Bergen (FCM Run 4); (3) The Ls Tromsø laboratory strain was established by crossing adults from the *Ls* Gulen strain with a partially outbred strain, *Ls* Oslofjord, originating from specimens collected in Oslofjord (Norway) and reared at the Aquaculture Research Station in Tromsø (FCM runs 1–3).

#### Collection of samples and tissue preparation

Newly hatched nauplii were obtained from gravid females, crushed in cold citrate buffer [75] containing 5% dimethyl sulfoxide (DMSO), filtered through a 30 μm nylon mesh and deep-frozen until use. Sperm and eggs were collected from the testes and the genital segment prior to fertilization, respectively, and the resulting samples briefly kept on ice prior to analysis. Somatic (cuticular and subcuticular) tissues obtained from the cephalothorax (Supplemental Materials and Methods Fig. 1) of adult wild or laboratory specimens were crushed, and treated in the same way as the newly hatched nauplii. Specimens were squashed onto slides according to Clower and co-workers [54] except that a freeze-cracking technique was added.

#### Flow cytometry analysis

Aliquots of target (sea lice) and internal reference (male human and/or chicken) cells were analyzed using Propidium Iodide (PI) as fluorescent stain following previously reported methods [76]. The mean DNA content of 5000 – 10,000 cells per sample was measured with a CyFlow^®^Ploidy Analyser equipped with a green laser.

Nuclear DNA contents of target species were estimated in relation to an assigned 2*C* value of 7.00 pgDNA/nucleus for human leukocytes and 2.50 pg DNA/nucleus for chicken erythrocytes [52] according to the formula:

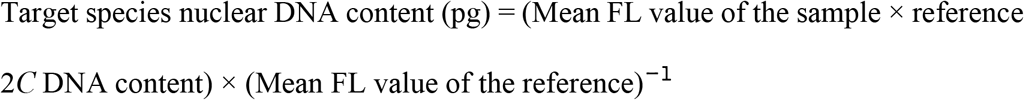

### Feulgen image analysis densitometry

#### Field and laboratory populations

Genome size measurements were obtained from each of three adults (females) of the *Ls* 1a laboratory strain described elsewhere [77], whose ovaries served as the source material of DNA used in the nanopore DNA sequencing and six adults (three males and three females) of *L. salmonis salmonis* from *Ls* Gulen laboratory strain used in the FCM studies. A single adult female was collected from the wild in Copscook Bay, Maine in 2018. Specimens were immediately preserved in undenatured >95% alcohol.

#### Feulgen staining and scanning microdensitometry

All slides were squashed and stained with Schiff reagent according to previously reported methods [45,54], with few modifications. Nuclei were measured using a Zeiss Axioscope A1 equipped with a 63X oil objective and a Qimaging Bioquant PVI CCD camera. Scanning microdensitometric software (Bioquant Image Analysis; Bioquant Life Sciences 2018 program) was used to determine the IODs of the nuclear DNA contents of individual somatic nuclei. We selected for measurement only nuclei that possessed a granular and slightly diffuse appearance and lacked visible pink background; these nuclei were found mostly at the perimeter or outside the carapace (Fig. 2c,d). Nuclei with relatively small areas and dense staining indicating DNA compaction (Fig. 2e) or very diffuse and large areas (Fig. 2f) are less likely to provide accurate measurements. The Bioquant software used to measure IODs has a conservative estimate of resolution of 0.5 pg DNA per nucleus according to the manufacturer. The mean IOD value of the hen was used to convert the IODs of each *L. salmonis salmonis* specimen to picograms, using the following equation:

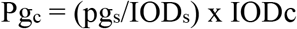

where pg_c_ is the unknown amount of pg DNA per nucleus of *L. salmonis salmonis* pg_s_, 2.5 pg is the amount of DNA in the standard hen nucleus, IOD_s_ is the average IOD value of the hen, and IODc is the IOD value of *L. salmonis*. Photographs were taken at 100X magnification with a Nikon Eclipse Ti-2 microscope equipped with a PlanApo objective (N.A. 1.45) and QImaging DS RI2 camera.

Reference standards for conversion of integrated optical density (IOD) units to picograms (pg) included mutant white eyed female *Drosophila melanogaster* (0.40 pg DNA per nucleus), erythrocytes of hen *Gallus domesticus* (2.5 pg DNA per nucleus) and trout *Onycorhynchus mykiss* (5.2 pg DNA per nucleus), and leucocytes of male human *Homo sapiens* (7.0 pg DNA per nucleus) whose values were based on previous works [78,79] and the Animal Genome Size Database [52]. The calibration curve computed for standards in the staining batch containing the *Ls*1a strain yielded an R^2^ = 0.997 (Supplementary Methods Fig. 2), indicating quantitative staining over a range of 0.40 – 7.0 pg DNA per nucleus. Only hen and trout standards (Fig. 2a,b) were used in the staining batch with *Ls* Gulen and Maine specimens.

The mean nuclear DNA contents are reported as 2*C* values in picograms (pg) and converted to gigabase (Gb) pairs (1 pg DNA = 0.978 Gb) for both FCM and FIAD derived estimates [80].

### Statistical analyses

Differences in nuclear DNA content of somatic tissues between LsTromsø nauplii (FCM Run 1 and 2) and *Ls* Gulen germinal (eggs and sperm) or somatic tissues of LsTromsø and *Ls* Gulen adult males and females (FCM Run 3–4), as well as those of *Ls* Gulen adults obtained using FIAD, were analyzed by Students *t*-test. Analysis of variance (ANOVA) was used to detect significant differences in nuclear DNA content of somatic tissue of adult wild caught and laboratory LsTromsø strain *L. salmonis salmonis* (FCM Run 5) using fluorescence (FL) PI values as dependent variable and gender and strain (laboratory or wild) as factors. In Exploratory Data Analysis (EDA), Grubbs’ test was used to detect presence of outliers and Levene’s and Shapiro-Wilk tests were used to test homogeneity of variances among groups. Statistical analyses were performed using IBM SPSS Statistics v.25 software. Differences were accepted as significant when P< 0.05. Data are reported as mean ± standard error (SE). The Shapiro-Wilks test applied to nuclei within each of 10 specimens measured using FIAD revealed no departures from normality. Differences between male and female genome sizes based on FIAD were tested using two sample, two tailed Student *t*-tests.

## Abbreviations

BUSCO: Benchmarking Universal Single-Copy Orthologs
FIAD: Feulgen image analysis densitometry
FCM: Flow cytometry
Fl: Fluorescence
Gbp: Giga base pairs
Mbp: Mega base pairs
PI: Propidium iodide
SNP: Single Nucleotide Polymorphism

## Acknowledgements

We thank James Madison University (JMU) Physics Department for constructing the liquid nitrogen table, Adrian Streit for methodological advice, Kris Kubow for imaging support and preparation of plate, Harrison Giknovorian for technical assistance with squashing and Feulgen reaction, Emilly Schutt for preparation of the graph, Ken Roth for supplying the human blood, Marquis Walker for providing the *Drosophila* and methods for histological preparation, and Michaёl Bekaert, Tyler Elliott, Ryan Gregory, Nick Jeffery, and Ben Koop for critical discussions. Work at JMU was supported by NIH 1R15GM104868 and NSF-DBI-1725855 to GAW and others, NSF-DEB 1948267 to GAW and a grant to GAW from the James Madison University Program of Grants for Faculty Assistance. The publication charges for this article have been funded by a grant to SP from the publication fund of UiT, The Arctic University of Norway. The authors are grateful to Anette Hustad, Linn Svendheim and Svenn Rune Hansen at the Aquaculture Research Station in Tromsø and to Sussie Dalvin (IMR, Bergen) for sea lice samples collection. We acknowledge the veterinarian Diogo Costa Ramos Da Rocha Marques for supplying the chicken blood and Goran Kauric at the University Hospital of North Norway (UNN, Tromsø) for processing the human blood samples.

## Author contributions

GAW, RSM and SP - conceptualized and designed the work. GAW and SP - conducted cytogenetic (FIAD, Flow Cytometry) analyses and data handling. RSM and KM - performed bioinformatic analyses. GAW, RSM, KM and SP - drafted and wrote the main manuscript. RP developed the freeze-cracking method for FIAD measurements. All authors reviewed the manuscript.

